# The P-selectin ligand PSGL-1 (CD162) is efficiently incorporated by primary HIV-1 isolates and can facilitate trans-infection

**DOI:** 10.1101/2021.06.29.450454

**Authors:** Jonathan Burnie, Arvin Tejnarine Persaud, Laxshaginee Thaya, Qingbo Liu, Huiyi Miao, Stephen Grabinsky, Vanessa Norouzi, Paolo Lusso, Vera A. Tang, Christina Guzzo

## Abstract

While P-selectin glycoprotein ligand-1 (PSGL-1/CD162) has been studied extensively for its role in mediating leukocyte rolling through interactions with its receptor, P-selectin, recently, it was identified as a novel HIV-1 host restriction factor. One key mechanism of HIV-1 restriction is the ability of PSGL-1 to be physically incorporated into the external viral envelope, which effectively reduces infectivity by blocking virus attachment through the steric hindrance caused by its large ectodomain. Importantly, a large portion of the literature demonstrating the antiviral activity of PSGL-1 has utilized viruses produced in transfected cells which express high levels of PSGL-1. However, herein we show that virion-incorporated PSGL-1 is far less abundant on the surface of viruses produced via infection of physiologically relevant models (T cell lines and primary cells) compared to transfection (overexpression) models. Unique to this study, we show that PSGL-1 is incorporated in a broad range of HIV-1 and SIV isolates, supporting the physiological relevance of this incorporation. We also report that high levels of virion-incorporated PSGL-1 are detectable in plasma from viremic HIV-1 infected individuals, further corroborating the clinical relevance of PSGL-1 in natural infection. Additionally, we show that PSGL-1 on viruses is functionally active and can bind its cognate receptor, P-selectin, and that virions captured via P-selectin can subsequently be transferred to HIV-permissive bystander cells in a model of trans-infection. Taken together, our data suggest that PSGL-1 may have diverse roles in the physiology of HIV-1 infection, not restricted to the current antiviral paradigm.

**IMPORTANCE:** PSGL-1 is an HIV-1 host restriction factor which reduces viral infectivity by physically incorporating into the envelope of virions. While the antiviral effects of PSGL-1 in viruses produced by transfection models is profound, HIV-1 continues to remain infectious when produced through natural infection, even when PSGL-1 is incorporated. To study this discordance, we compared the differences in infectivity and PSGL-1 abundance in viruses produced by transfection or infection. Viruses produced via transfection contained unnaturally high levels of incorporated PSGL-1 compared to viruses from primary cells, and were much less infectious. We also found PSGL-1 to be present on a broad range of HIV-1 isolates, including those found in plasma from HIV-infected patients. Remarkably, we show that virion-incorporated PSGL-1 facilitates virus capture and transfer to HIV-permissive host cells via binding to P-selectin. These findings suggest that PSGL-1 may also work to enhance infection *in vivo*.

## INTRODUCTION

P-selectin glycoprotein ligand-1 (PSGL-1/CD162) is an adhesion molecule expressed on all leukocytes that plays a critical role in the early stages of inflammation due to its ability to bind P-, E- and L- selectins (1–5). In particular, PSGL-1 has been well-characterized for its involvement in leukocyte recruitement (rolling and tethering) and extravasation into tissues (reviewed in (1, 3, 4, 6, 7)). Stucturally, PSGL-1 is a highly glycosylated homodimeric transmembrane protein, with an extracellular domain (ECD) of 50-60 nm in length that extends far out from the cellular surface (1, 8, 9). In 2019, Liu *et al.* were the first to identify PSGL-1 as a novel HIV restriction factor in activated human CD4+ T cells, with inherent antiviral activity through a variety of mechansims in host cells (10). Notably, using viruses produced through transfection, Liu *et al.* showed that when increasing amounts of PSGL-1 plasmid DNA (pDNA) were co-transfected with an HIV-1 infectious molecular clone (IMC), virion infectivity was reduced in a dose-dependent manner (10). Upon further investigation virions were also shown to physically incorporate PSGL-1 using electron microscopy and Western blotting techniques, and this virion incorporation of PSGL-1 was identified as an additional mechanism to diminish HIV-1 infectivity (10). Experiments performed by Murakami *et al*. and Fu *et al.* in 2020 also demonstrated through knock-down of PSGL-1 in Jurkat cells and CRISPR-mediated KO of PSGL-1 in primary cells respectively, that physiological levels of PSGL-1 on the cell surface can also result in a modest impairment of particle infectivity (11, 12).

Subsequent studies examining the antiviral effects of virion-incorporated PSGL-1 demonstrated that PGSL-1 could block the binding of virions to target cells (11, 12). More specifically, it was shown that the large ECD of PSGL-1 was required for this inhibitory effect and that PSGL-1 also has a broad spectrum antiviral effect when ectopically expressed in the envelopes of other viruses, including murine leukemia virus, influenza A and SARS CoV-1 and −2 (11, 13, 14). Additional characterization of virion-incorporated PSGL-1 revealed a concomitant reduction in the incorporation of the HIV envelope glycoprotein (Env) into virions by sequestering gp41 at the cell membrane (15), providing another measure by which PSGL-1 can restrict virion infectivity. Research on other structurally similar proteins with large extracellular domains has also shown similar inhibitory effects on viral infection and in line with this concept, PSGL-1 was recently reported to be a part of a larger group of antiviral proteins termed Surface-Hinged, Rigidly-Extended Killer (SHREK) proteins (14).

While much has been rapidly uncovered about the antiviral functions of PSGL-1 on HIV-1 and other viruses since its identification as a host restriction factor in 2019 (10), many unanswered questions about the function of PSGL-1 in the physiology of HIV-1 infection remain. Indeed, while the initial proteomic screen that identified PSGL-1 as a host restriction factor and limited others employed activated primary CD4+ T cells (10–12), the majority of experiments performed to characterize the antiviral functions of this protein have used viruses produced via co-transfection of PSGL-1 and viral constructs in adherent cell lines. While transfection models are commonly used to produce viruses and express specific host proteins, it is well recognized that the cell type used to produce viruses can alter the composition of cellular proteins incorporated into the virion surface (16–18). Importantly, when transfecting DNA, the levels of gene expression can be highly variable and can be up to 100-fold higher than natural expression, since gene expression is greatly dependent on the strength of the promoter used (19–21). It is also important to note that PSGL-1 functionality is strongly dependent on the cell type and activation state of the producer cells since it requires highly specific glycosylation patterns to bind selectins (3, 4, 7, 22, 23). While many antiviral properties of PSGL-1 have been characterized to date, it remains unknown whether virion-incorporated PSGL-1 can bind its cognate receptor, P-selectin. This additional binding capacity may impact HIV-1 pathogenesis and provide additional roles for virion-incorporated PSGL-1 *in vivo,* beyond its current antiviral classification. Indeed, many cellular proteins within the HIV-1 envelope are known to retain their biological function, which can enhance viral infectivity and modify pathogenesis (16, 24–27). Furthermore, no studies to date have shown if PSGL-1 is present on circulating strains of viruses in HIV-1 infected patients, which is critical for establishing the clinical relevance of PSGL-1 in HIV-1 infection.

Herein we show that levels of PSGL-1 on virus preparations produced from cells transfected to express PSGL-1 are not representative of the level of PSGL-1 present on primary viral isolates. Additionally, we demonstrate with novel techniques that while PSGL-1 can inhibit the infectivity of transfected viruses in a dose-dependent manner, viruses produced in primary cells remain infectious despite endogenous levels of PSGL-1 incorporation, as detected through antibody-based virus capture assays and flow virometry. Furthermore, we are the first to show that PSGL-1 is present on a wide range of HIV-1 isolates produced in primary peripheral blood mononuclear cell (PBMC) cultures and on circulating virions in the plasma of HIV-1 infected patients at various stages of disease progression and viremia. Notably, we found that HIV-1 Env (gp120) remains present at detectable levels on all of the primary isolates cultured *in vitro*, despite PSGL-1’s ability to reduce levels of Env on virions (11, 15). Most importantly, we demonstrate that viruses displaying PSGL-1 on their surface can be captured with P-selectin (CD62P) and transferred to permissive cells in a model of trans-infection. These data suggest novel roles for PSGL-1 in fueling HIV-1 infection and pathogenesis, that further extends the functionality of virion-incorporated PSGL-1 beyond the previously described antiviral activity.

## RESULTS

### Overexpression of PSGL-1 in HEK293 cells markedly reduces infectivity of progeny virions, while viruses produced by natural infection of T cells and PBMC retain infectivity

In our recent work we phenotyped pseudoviruses engineered to display the virion-incorporated cellular proteins integrin α4β7, CD14 and PSGL-1 (CD162). This work led us to the striking observation that PSGL-1 was detected at markedly higher levels on virion surfaces compared to other host proteins, despite all pseudoviruses (PV) being produced with equal amounts of pDNA for host protein expression (28). Since we observed PSGL-1 to be incorporated to a greater extent than other host proteins on the surface of pseudoviruses, we wanted to determine how phenotypically similar pseudoviruses containing PSGL-1 were to viruses produced in more physiologically relevant model systems, such as infection of T cell lines and primary cells. To this end, we first produced two virus models via transfection: PV through co-transfection of plasmids expressing an HIV-1 backbone and envelope (SG3^ΔEnv^ backbone + BaL.01 envelope), and full-length viruses with an NL4-3 infectious molecular clone (IMC). In both virus types (PV and IMC), in addition to HIV expression plasmids, we also co-transfected different amounts of PSGL-1 pDNA to explore whether we could generate virus progeny with low, medium and high levels of PSGL-1 in the HIV-1 envelope (designated PSGL-1^Low^, PSGL-1^Med^, PSGL-1^High^ respectively; Table S1). Of note, PSGL-1^High^ virions were created using pDNA amounts that we typically employ to generate viruses with host proteins, followed by step-wise reductions to generate medium and low levels of expression. As a control, viruses without PSGL-1 (PSGL-1^Neg^) were engineered with HIV-1 pDNA alone. For comparison to our transfected viruses, we used T cell lines or primary PBMC to produce virus by *in vitro* infections. For the former model, we infected the H9 and Jurkat T cell lines with the HIV-1 IIIB isolate. Primary PBMC were infected with the HIV-1 isolates IIIB, BaL, and an NL4-3 IMC (previously harvested from transfected HEK293 cultures), which were all minimally passaged in PBMC. We chose the lab-adapted HIV-1 isolates IIIB (X4-tropic) and BaL (R5-tropic) for ease of generating high levels of infection in these cell types. Infectivity of the virus stocks was tested in the commonly used TZM-bl reporter cell line using luminescence as a readout for infection (12, 29–31). To ensure that we could accurately compare the differences in infectivity observed with differential amounts of virion-incorporated PSGL-1, TZM-bl cells were infected with serial dilutions of virus stocks with input normalized by p24 across each independent virus cluster (i.e., within the group of PVs, IMCs, and T cell line viruses; Fig. 1A-C). Since three different PBMC donors were used to generate the PBMC viral stocks, more variation was present in the viral titres, due to the nature of donor variability. PBMC viruses were tested at their undiluted concentration to ensure the range of variability in infectivity seen in isolates produced in primary cells could be visualized (Fig. 1D). Importantly, analyses of the role of PSGL-1 in virus infectivity were only made within a virus cluster (i.e. within the group of PVs, IMCs, etc.) and not between the model systems used for virus production.

**FIG 1.**
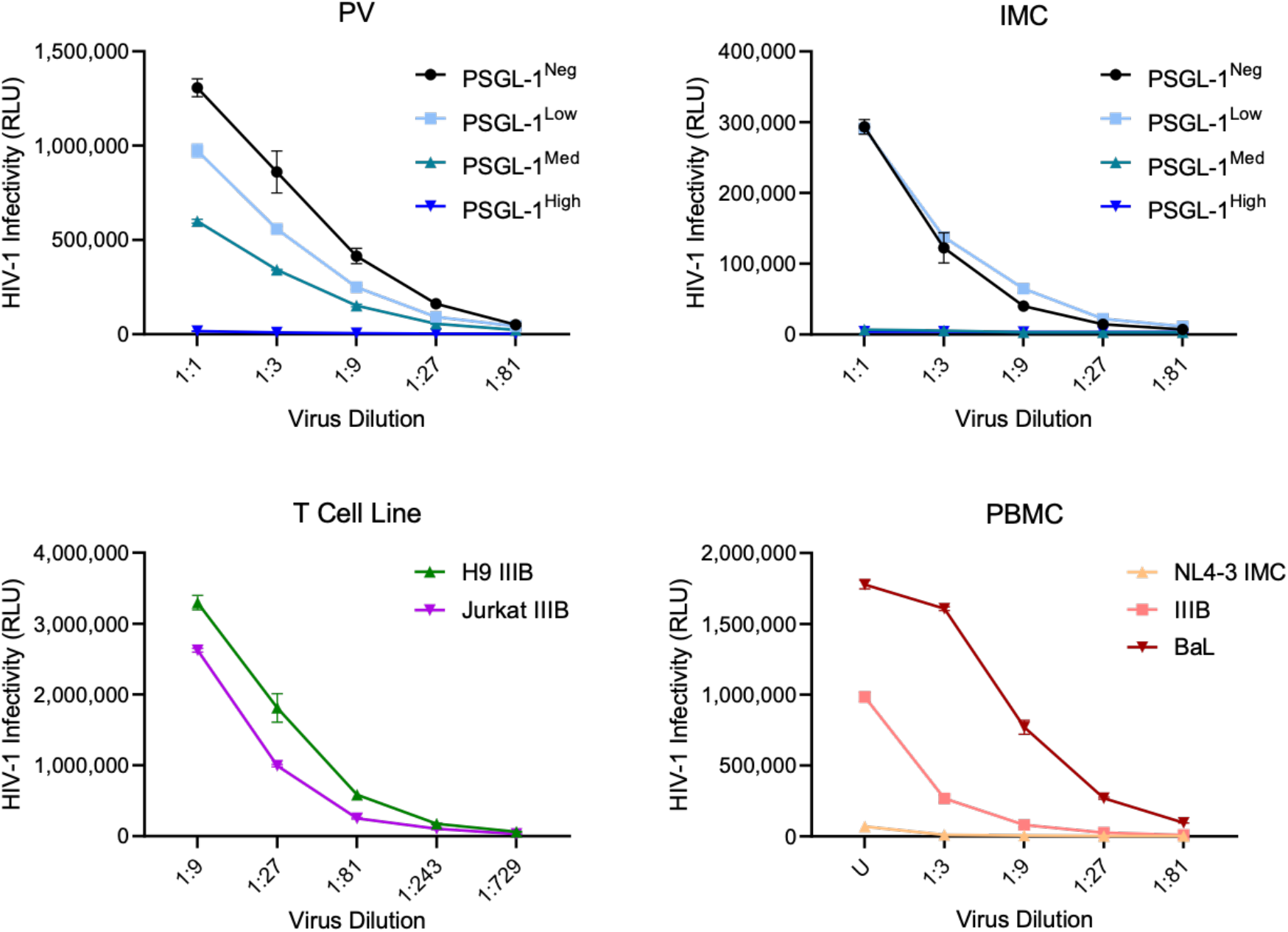
Infectivity of virus stocks produced via transfection versus infection. (A) Pseudoviruses (PV) produced through the co-transfection of the viral backbone SG3^ΔEnv^ HIV-1 (2 µg) with varying levels of BaL.01 envelope (0.75 - 1 µg) and PSGL-1 (2.5-250 ng) plasmids, as outlined in Table S1, were normalized based on viral p24 (displayed as 1:1 in graph) and were subjected to 3-fold serial dilutions. Control (PSGL-1^Neg^) virus produced without PSGL-1 plasmid DNA was assayed in parallel. All diluted virus stocks were incubated with TZM-bl reporter cells for 48 hours before infectivity was measured using luminescence (relative light units; RLU). (B) Full-length replication-competent viruses either devoid of or containing various levels of PSGL-1 were produced through transfection of the infectious molecular clone (IMC) pNL4-3 alone (2 µg) or with the IMC and PSGL-1 plasmid DNA (2.5 ng - 1 µg) respectively, as outlined in Table S1. A negative control vector (empty vector) was used to normalize the total amount of plasmid used for transfection to 3 µg for the production of all IMC viruses. Viruses were diluted and tested as in (A). (C) Replication-competent HIV-1 IIIB was passaged in the H9 and Jurkat T cell lines and assayed for infectivity using TZM-bl cells as in (A&B). Data points from higher virus dilutions (1:243 and 1:729) are shown to display the full titration of infectivity. (D) The lab-adapted HIV-1 isolates BaL and IIIB produced in PBMC, and NL4-3 IMC produced through HEK293 transfection were all passaged once in activated PBMC. The resulting progeny virions were harvested and tested undiluted (U) or at the indicated dilution on TZM-bl cells, as described above. Results are displayed as mean +/-standard deviation of samples tested in duplicate.

As expected, the infectivity of all transfected viruses was greatly diminished when PSGL-1 was present at high levels within virions (Fig. 1A and 1B; PSGL-1^High^). The dose-dependent potency of PSGL-1 as an antiviral factor was most evident in the pseudoviruses, which showed a clear stepwise decline in infectivity as increasing amounts of PSGL-1 pDNA were used to produce virions (Fig. 1A). Remarkably, even the PSGL-1^Low^ PVs (which were created with very low levels of pDNA, (Table S1) displayed reduced infectivity compared to PSGL-1^Neg^ PVs (Fig 1A). Interestingly, no defined differences in infectivity were present between the IMC PSGL-1^Med^ and PSGL-1^High^ viruses, despite the 4-fold difference in PSGL-1 pDNA used to produce these two virus types (Table S1). Most strikingly, the infectivity of viruses produced in T cell lines and PBMC was considerably higher than transfected viruses (PVs and IMCs) with PSGL-1 (> 1 million RLU; Fig. 1C and 1D) except for one PBMC isolate (NL4-3 IMC; Fig. 1D). This exception was likely due to the fact that passaging of infectious molecular clones *in vitro* can alter the viral phenotype, including infectivity (32, 33). However, since all of the viruses were produced in different model systems, we focused comparisons of infectivity within each respective virus cluster and not across the different virus production models (i.e., PV, IMC, etc.). Indeed, prior work has shown that pseudovirus and full-length IMC viruses cloned from the same viral isolates could still display vastly different levels of infectivity depending on transfection conditions (34). Regardless, since the impact of PSGL-1 on viral infectivity was remarkably clear in transfected virus stocks, but not those produced via natural infection, we proceeded to compare the relative levels of PSGL-1 on the surface of these viruses.

### Virion-incorporated PSGL-1 is more abundant on viruses produced in cells transfected to express PSGL-1 versus those produced via infection of primary cells

To semi-quantitatively assess the amount of PSGL-1 in the envelopes of our different virus preparations, we used a previously described immunomagnetic virion capture assay (24, 28, 35, 36). This technique was chosen for its advantages over Western blot as it allows us to selectively assay for proteins that are found only on the viral surface (14, 37–39). To compare the relative amounts of PSGL-1 and gp120 present on our range of viruses, we performed antibody-mediated virion capture on normalized virus inputs (normalized p24 input across all viruses tested) with an anti-PSGL-1 monoclonal antibody (mAb) versus an anti-gp120 mAb (Fig. 2). Through the use of normalized virus inputs across all capture reactions, direct comparisons between the amount of antibody-mediated capture can reflect the relative levels of virion incorporated PSGL-1 and gp120. As expected, the anti-PSGL-1 antibody captured all of the transfected virus preparations engineered to contain PSGL-1, while no PSGL-1 capture was present with control viruses (PSGL-1^Neg^) devoid of PSGL-1 (Fig. 2A and 2B). Stepwise differences in levels of PSGL-1 capture were seen in the PSGL-1-positive (PSGL-1+) pseudoviruses based on their engineered designations (low, medium, high; Fig. 2A). Distinct differences in the amounts of PSGL-1 capture were also present in the PSGL-1+ IMC viruses, although no difference was apparent between the PSGL-1^Med^ and PSGL-1^High^ viruses, similar to what was seen in viral titration (Fig. 1B). Importantly, levels of gp120 capture were either very low or below background (as determined by capture with a non-specific antibody) on all of the PSGL-1+ pseudoviruses and the PSGL-1^Med^ and PSGL-1^High^ IMC viruses (Fig. 2A and 2B). While it may appear that some of these viruses are completely devoid of gp120 from the results of this assay, data from infectivity assays (Fig. 1) show that a minimal level of Env must be present within the infectious viruses. Furthermore, it is likely that the level of gp120 capture in these viruses is close to or at the threshold of detection for this type of capture assay. Indeed, gp120 has been reported to be present at low levels on circulating virions (8-14 spikes) (40) and PSGL-1 is also known to disrupt Env incorporation into virions (11, 15), so it is unsurprising that gp120 is less abundant on viruses produced from cells transfected to express PSGL-1. Moreover, since the viral accessory protein Vpu, which works to counteract this sequestration, is expressed in the late stages of viral replication (41), we hypothesized that transfected viruses generated with a short (2 day) production protocol would be more impacted by Env sequestration than viruses produced in lengthier (7-14 day) infection protocols. In line with this, anti-PSGL-1 capture on viruses produced in T cell lines and PBMC demonstrated that PSGL-1 was present on all of the isolates tested (Fig. 2C and 2D). Interestingly, levels of PSGL-1 capture were much lower in these more physiologically relevant viruses than viruses produced by transfection (up to 30-fold less virion capture). It is also important to note that gp120 was detectable above background in all of the T cell line and PBMC viruses tested, which may be related to why most of these viruses were more infectious than the PSGL-1+ transfected viruses. While these results confirmed that there were differences between the levels of PSGL-1 and gp120 capture on viruses produced in different cell types, antibody capture assays are only semi-quantitative and report on the average phenotype of the sample. Hence, similar to other bulk techniques such as Western blot, this assay lacks the resolution to interrogate a heterogenous sample containing phenotypically distinct virions with individual variation in the amounts of host and viral proteins. To gain more quantitative data on the abundance of PSGL-1 on individual virions, we decided to stain virus particles and analyze by flow virometry, for its advantages in providing high throughput, single virion analysis in a calibrated and quantitative readout (42–45).

**FIG 2.**
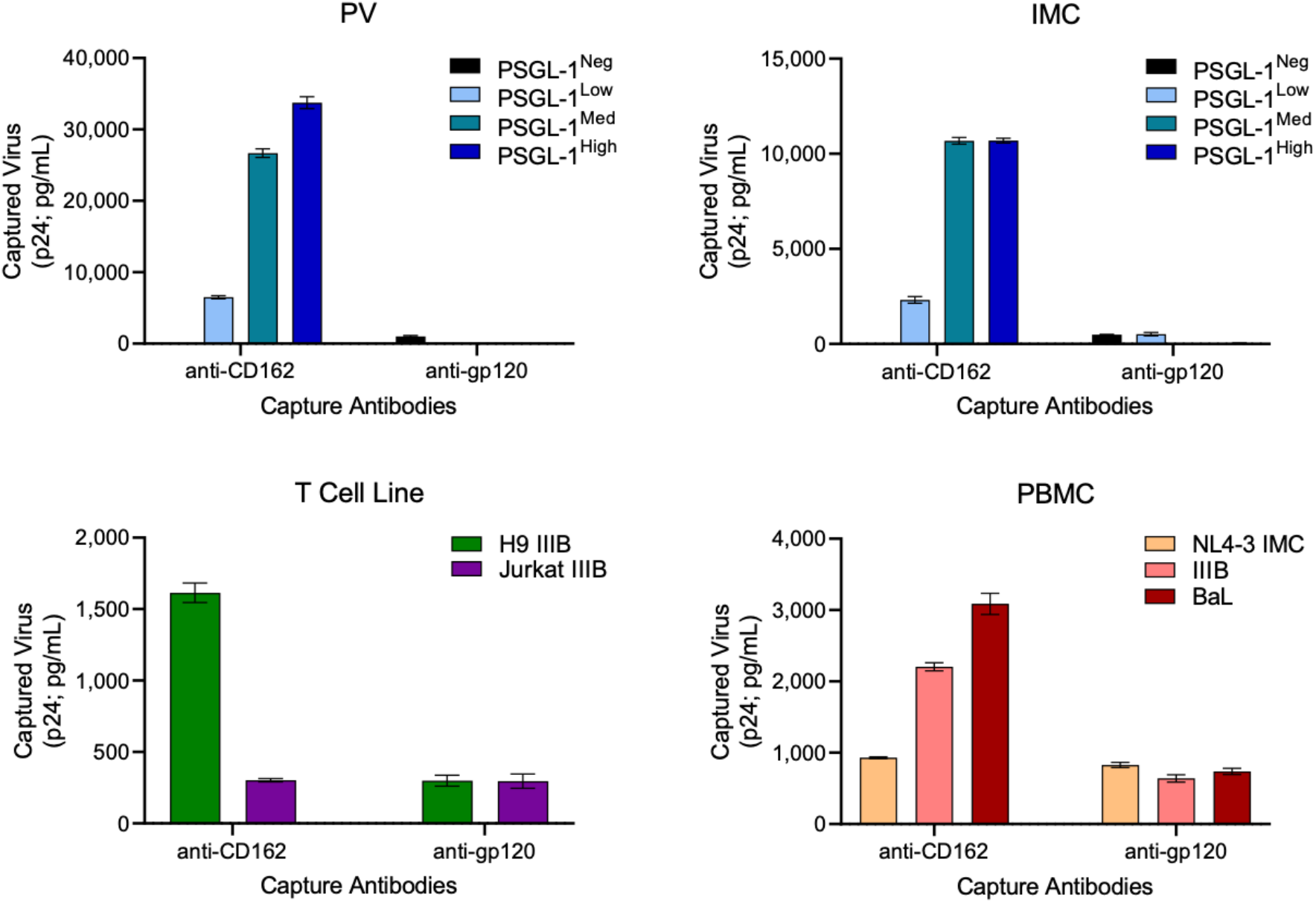
Semi-quantitative comparisons of virion-incorporated PSGL-1 and gp120 on virus stocks produced via transfection versus infection. (A) Virion capture assays were performed with immunomagnetic beads armed with anti-PSGL-1 (clone KPL-1) or anti-gp120 (clone 2G12) with normalized inputs (5.25 ng of p24 per condition) of pseudovirus (PV), (B) IMC viruses, (C) T cell line viruses, and (D) PBMC viruses (as described in Fig. 1). Bead-associated virus was lysed and HIV-1 p24_Gag_ was quantified using p24 AlphaLISA as an indicator of virus capture. Results are displayed as mean +/-standard deviation of samples tested in duplicate. Levels of background capture as detected using an isotype control antibody were subtracted from the displayed values.

### T cell and PBMC viruses incorporate lower amounts of PSGL-1 than viruses produced in cells transfected to express PSGL-1

We have previously provided detailed methodology showing how HIV-1 can be detected by light scatter on sufficiently sensitive cytometers (28, 46), and that virion-incorporated host proteins (including PSGL-1) can be quantified in the HIV-1 envelope of pseudoviruses using flow virometry (28). More specifically, by using fluorescent and light scatter calibration reference materials along with calibration software, arbitrary light intensities from stained virus samples can be calibrated and expressed in standard units allowing for quantification and comparison of proteins on virions. These standardization methods allow us to draw close estimates of the total number of antibodies bound to each virion, which can be used as a proxy for total number of proteins on individual viral particles (28). Of note, in line with the scope of this study, we focused our efforts on PSGL-1 staining and quantitation, and not HIV-1 Env, since it is currently below our threshold for detection with conventional fluorescent labelling used in our staining protocol. Before staining our viruses for acquisition, we validated that we could discriminate virus from background instrument noise that is present from running cell culture media alone through the cytometer (Fig. S1).

Next, we performed labelling of pseudoviruses and IMCs that we knew contained high levels of PSGL-1 in the envelope with a PE-labelled anti-PSGL-1 mAb (same antibody clone used in virion capture assays). We observed stepwise increasing amounts of median PSGL-1 PE labelling (as reported in calibrated units of molecules of equivalent soluble fluorophore; MESF) on both sets of transfected viruses (PV and IMC; Fig. 3A). The median MESF values are presented with robust standard deviation (SD), since this SD calculation is less skewed than the conventional SD by outlying events that could be caused by background non-virus events. To determine MESF values, viruses were first gated by side scatter (Fig. 3) and then a secondary gate spanning the same SSC profile (not shown) was placed on the population of viruses that were above the level of background fluorescence (10 MESF) to generate MESF and SD statistics. This ensured that only viruses that were successfully labelled were contributing to our MESF counts. The pseudoviruses showed the best dynamic range of successful staining, with MESF values as 15 ± 5 SD, 35 ± 45 SD and 145 ± 120 SD (Fig. 3A, top row), for the PSGL-1-Low, - Med and -High viruses, respectively. Despite the use of robust instead of conventional standard deviation, a range of error up to 1.3-fold the reported median was still present since the virus gates that were used (Fig. 3) were set to span a wide range of PE MESF values. This broad gate can contain low levels of coincident events and antibody background fluorescence that contribute to the SD that are not removed since our virus labelling protocol does not include wash steps. We chose to accept this level of error since using this broad SSC gate allowed for the most accurate comparison of all of the different virus clusters tested.

**FIG 3.**
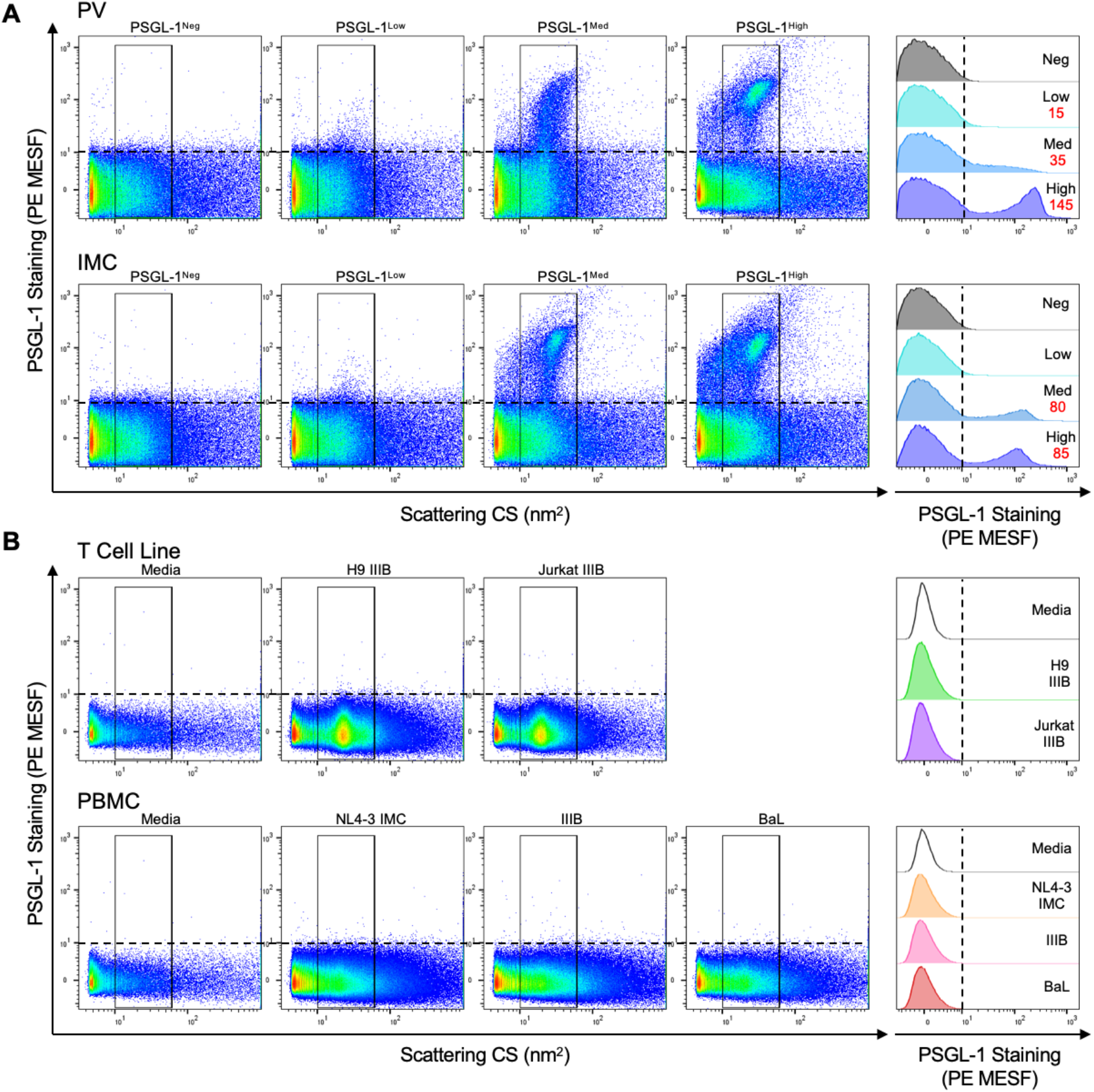
Staining and quantitation of virion-incorporated PSGL-1 using flow virometry. (A) Staining of transfected pseudoviruses (PV) and full-length viruses (IMC) with different PSGL-1 phenotypes (negative, low, medium, high) with a primary PE-conjugated anti-PSGL-1 antibody overnight at 4 °C. A comparison of PSGL-1 staining on all virus populations with the median PE MESF value of each population (values in red) shown in the rightmost panel for each virus type (PV, IMC). PSGL-1 staining that yielded MESF values below 10 are not displayed. MESF values were determined from the population of gated viruses that appear above the horizontal dotted line on virus dot plots which indicates background fluorescence and the limit of instrument detection (10 MESF). An equivalent vertical line depicted on each histogram denotes the same level of background fluorescence. (B) Staining of viruses produced in T cell lines (upper panel) and primary cells (PBMC; lower panel) displayed as in (A). Viruses were acquired on the cytometer for 2 minutes on low. Data shown are representative of two technical replicates. Virus staining shown in (A) uses 0.8 µg/mL of antibody while staining in B uses 0.1 µg/mL. These data were selected from a four-point antibody titration (as shown in Fig. S2) since they showed optimal signal and minimal background fluorescence from unbound antibody based on the conditions tested.

A small difference in the median PE MESF values for the PSGL-1^Med^ and PSGL-1^High^ IMC viruses (80 ± 70 SD and 85 ± 65 SD) was observed (Fig. 3A, bottom row), but the standard deviation for the two conditions overlapped similar to what was seen in viral titration (Fig. 1) and in virus capture (Fig. 2). With respect to the PSGL-1^Low^ IMC virus, while more labelling was apparent on the stained virus samples than the cell culture media (Fig. 3A), no measurable labelling was present on the dot plots above the background fluorescence (horizontal dotted line; Fig. 3A), and the median PE MESF value for the PSGL-1^Low^ IMC virus fell within the range of background for the cytometer (10 PE MESF). As expected, control viruses that were devoid of PSGL-1 (PSGL-1^Neg^) also did not exhibit positive staining above background fluorescence. After validating that our flow virometry protocol was effective and sensitive enough to detect differences in surface levels of virion-incorporated PSGL-1 in transfected viruses, we next acquired our T cell line and PBMC viruses (Fig. 3B). We observed that viruses produced in T cell lines produced more homogenous virus populations than those seen in viruses made in PBMC, as expected given the nature of the culture conditions. Additionally, T cell line viruses were more monodisperse than viruses propagated in PBMC and produced ‘cleaner’ dot plots with less background attributable to non-virus events. The heterogeneity in PBMC virus scatterplots compared to those of T cell line viruses is likely attributable to differences in extracellular vesicles produced by cell lines versus PBMC cultures (Fig. 3B upper row vs bottom row). Surprisingly, while very low levels of PSGL-1 staining are apparent by eye in T cell line and PBMC viruses (Fig. 3B), this minute level of staining also fell within the range of background fluorescence on the cytometer when reported in standardized MESF units.

Since the dim fluorescence from low abundance antigens expressed on small particles may be masked by background auto-fluorescence or be in the range of instrument noise (47, 48), reducing background fluorescence and optimizing signal to noise ratio are critical in flow virometry. To ensure that we were not underestimating the levels of PSGL-1 staining due to insufficient antibody concentration, we tested four different antibody concentrations on our virus preparations, but were still unable to see high levels of staining on T cell line or PBMC viruses (Fig. S2). Since we were able to detect PSGL-1 on PBMC viruses using bead-based capture (Fig. 2D), we anticipated that PSGL-1 on primary virions were at levels that were simply too low to detect using our current flow virometry protocols. Indeed, since our capture assays showed that the amount of PSGL-1 on virions produced in T cell lines and PBMC was similar in magnitude to that of gp120 on the viruses tested (i.e. < 3-fold difference; Fig. 2C and 2D), and since we know that anti-gp120 labelling is currently at the cusp of our detection sensitivity for our flow cytometer (roughly ∼10 PE MESF which is approximately equivalent to ∼10 antibodies bound to each virus) (28), these data are in line with the limit of detection that we would expect using this method.

Notably, we were able to stain all of our transfected viruses and observe fluorescence that could be reported in MESF units well above the background for the majority of viruses. Thus, while we were unable to accurately report quantitation of PSGL-1 on primary viruses using this method, MESF calculations on transfected viruses demonstrated that flow virometry is robust for detection of molecules that are present within the working range of instrument detection. Indeed, we were also able to see successful labelling of PSGL-1 on transfected virus with an unlabelled PSGL-1 antibody paired with a fluor-labelled secondary antibody (Fig. S3). However, we only tested this on a small set of samples and did not pursue this method of staining further since the introduction of a secondary antibody reduced the fluorescence signal.

### PSGL-1 is incorporated by a broad range of HIV-1 and SIV isolates and is present on virions in plasma from HIV-infected patients

After demonstrating that there were large differences in the levels of PSGL-1 present on virions produced in different cell types, we next wanted to determine how abundant PSGL-1 was on a broader range of HIV-1 isolates grown in primary PBMC and in clinical samples. To determine the breadth of PSGL-1 virion-incorporation, we used immunomagnetic virion capture as above to compare PSGL-1 and gp120 incorporation among a panel of lab-adapted and clinical HIV-1 isolates representing different co-receptor usage phenotypes and clades (Table 1), as we have performed previously for virion incorporated integrin α4β7 (24). The different viral isolates were tested undiluted in capture assays to determine the potential for PSGL-1 incorporation at a broad range of viral titres. The PSGL-1 antibody successfully captured all strains of HIV-1 tested, and in most cases at higher levels than anti-gp120 capture. Furthermore, the levels of PSGL-1 capture were independent of HIV-1 clade or coreceptor usage. We compared the relative amount of virion incorporated PSGL-1 to gp120 among the different viral isolates by calculating the ratio of virion capture with anti-PSGL-1 mAb to anti-gp120 mAb (Table 1, capture ratio). Across most isolates tested we observed an appreciable excess of virion incorporated PSGL-1 relative to gp120, with a ratio average of 4.6 among the 14 viruses tested. In contrast, the ratio on transfected PSGL-1^High^ IMC viruses was greater than 250 (Fig. 2), highlighting several orders of magnitude difference in these model systems for PSGL-1 incorporation. Importantly, the ratio could not be calculated for PSGL-1+ pseudoviruses since gp120 was undetectable using antibodycapture when PSGL-1 was present on pseudoviruses. Notably, the PSGL-1^Low^ IMC virus had a ratio that was more similar to that seen with primary viruses, demonstrating that physiological levels of PSGL-1 incorporation can be displayed on viruses produced through transfection when using very low levels of pDNA (Table S1).

**Table 1.** Virion incorporation of PSGL-1 and gp120 in a panel of clinical and laboratory HIV-1 and SIV isolates grown in primary human PBMC.

| Virus | Isolate | Clade | Co-receptor Usage | Input virus (pg) | Captured Virus |  | Capture Ratio (PSGL-1:gp120) |
| --- | --- | --- | --- | --- | --- | --- | --- |
|  |  |  |  |  | PSGL-1 (pg/ml) | gp120 (pg/ml) |  |
| HIV-1 | 92UG037 | A | R5 | 3113 | 337 | 237 | 1.4 |
| HIV-1 | BaL | B | R5 | 3793 | 2857 | 1404 | 2.0 |
| HIV-1 | ADA | B | R5 | 8544 | 6945 | 2808 | 2.5 |
| HIV-1 | JRFL | B | R5 | 565 | 314 | 60 | 5.2 |
| HIV-1 | IIIB | B | X4 | 7068 | 2485 | 5585 | 0.4 |
| HIV-1 | 07USLD | B | X4 | 6492 | 1414 | 1465 | 1.0 |
| HIV-1 | 07USLR | B | X4 | 6035 | 945 | 1215 | 0.8 |
| HIV-1 | 92HT599 | B | R5X4 | 2101 | 1382 | 555 | 2.5 |
| HIV-1 | 93UG065 | D | X4 | 7577 | 5732 | 4228 | 1.4 |
| HIV-1 | 93TH057 | E | R5 | 4642 | 2046 | 370 | 5.5 |
| HIV-1 | CMU06 | E | X4 | 4026 | 1241 | 173 | 7.2 |
| HIV-1 | 97BR019 | F | R5X4 | 9629 | 2976 | 170 | 17.5 |
| SIV | smE660.307 | - | R5 | 18035 | 6312 | 1071 | 5.9 |
| SIV | mac251.745 | - | R5 | 13044 | 4707 | 418 | 11.3 |

\* The level of virion incorporation was determined by measuring the amount of captured virus (via p24<sub>Gag</sub> for HIV-1 or p27<sub>Gag</sub> for SIV) by immunomagnetic beads armed with anti-PSGL-1 or anti-gp120 (2G12) monoclonal antibodies. All clinical isolates were minimally passaged *in vitro*.

Notably, we also detected the presence of PSGL-1 at high levels in two SIV isolates (Table 1), providing further support to the notion that PSGL-1 may be a broad-spectrum host restriction factor (11, 13, 14). Despite the ability for PSGL-1 to sequester gp41 and to disrupt envelope trimer incorporation into virions (11, 15), all of the HIV-1 and SIV isolates tested displayed appreciable levels of gp120 capture, above background as determined with capture using an isotype-matched antibody as a negative control. This may suggest that the viral accessory proteins in primary isolates may be sufficient to counteract the inhibitory effects generated by virion-incorporated PSGL-1 and permit higher levels of gp120 incorporation.

After determining that virion incorporated PSGL-1 was present on all of the virus stocks produced *in vitro*, we sought to determine whether PSGL-1 could be identified on virions that circulate in viremic HIV-infected individuals, as to our knowledge, this has yet to be described in the literature. To this end, we assayed virions in plasma samples from 12 patients at variable stages of HIV infection (acute/early to chronic) using immunomagnetic virus capture (Fig. 4). For this test, we used undiluted virus samples to allow us to observe the range of capture efficiency based on variable viremia levels (documented in Table S2). To increase the sensitivity of the assay and to enhance the success of virion capture from patient plasma samples which contain many inherent factors that can hinder this assay (e.g. plasma proteins, extracellular vesicles, etc.), we employed a slightly modified capture assay using biotin-conjugated antibodies and compatible microbeads which have shown enhanced levels of virion capture in our hands over the conventional protein-G Dynabead captures used earlier in this study. Among the 12 patient plasmas tested, all patients harbored virus with incorporated PSGL-1, although with variable efficiency of incorporation ranging from 16% to 70% of virus input (Fig. 4). As an additional control, we also assayed CD44 incorporation in the same plasma samples, since CD44 is known to be incorporated with high efficiency into HIV-1 virions (49–51). CD44 was also incorporated into virions from all patient plasmas tested, with a wide range of virus input recovered by capture with anti-CD44 (8-87%), indicating that PSGL-1 falls into a category of host proteins that are incorporated with high efficiency into HIV virions. Given the range of patient sample variability, the percentage of input virus captured for each respective condition is documented in Table S3. Interestingly, PSGL-1 was present in virions from both the acute/early and chronic stages of infection; however, the sample size tested here is not large enough to draw statistically significant conclusions about PSGL-1 incorporation at different stages of infection.

**FIG 4.**
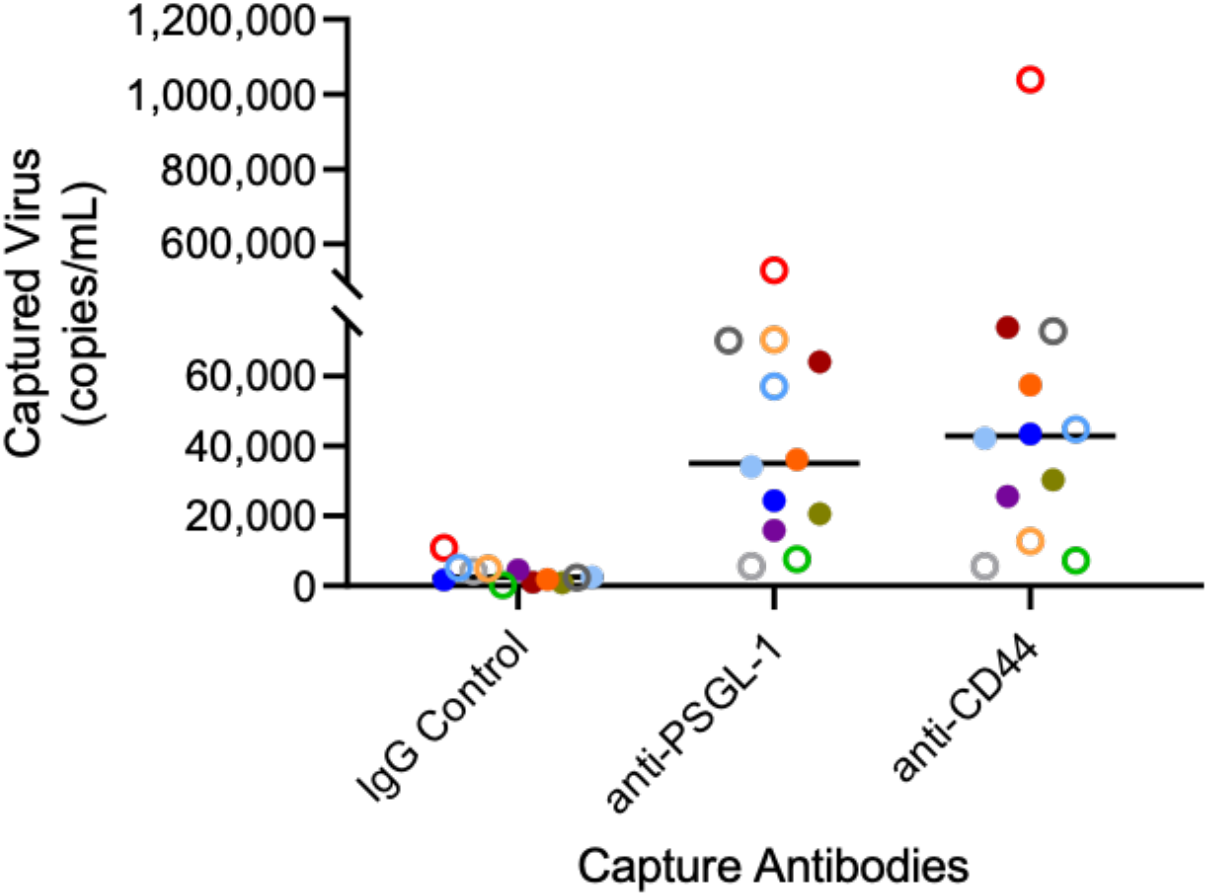
Virions circulating *in vivo* in HIV-infected patients contain PSGL-1 and CD44. Plasma samples from viremic patients ranging from early/acute to chronic stages of HIV-1 infection were tested in virion capture assays using an isotype control antibody (IgG control), anti-PSGL-1 or anti-CD44. Captured virus was lysed, followed by RNA extraction and quantitative real-time PCR for the detection of HIV-1 genome equivalents (in RNA copies/mL). The sample median for each antibody capture condition is displayed. Each unique symbol represents a different patient, with open circles denoting patients in the acute/early stage infection and filled circles as patients in the chronic stage of infection. Additional details regarding patient characteristics and the percentage of virion capture can be found in Table S2 and Table S3, respectively.

### Viruses with incorporated PSGL-1 can be captured by P-selectin and subsequently transferred to target cells for HIV-1 trans-infection

After quantifying virion-incorporated PSGL-1 on a variety of virus types and confirming that PSGL-1 was present on viruses from clinical samples, we next decided to assess whether virion-incorporated PSGL-1 was able to maintain its biological function and bind its cognate receptor, P-selectin. We speculated that this was highly plausible, since several other virion-incorporated proteins have been reported to maintain their physiological functions (16, 17, 24, 37, 39, 52, 53). To begin investigating this, we tested the capacity of transfected IMC viruses with and without PSGL-1 (PSGL-1^High^ and PSGL-1^Neg^) to bind recombinant P-selectin immobilized on plastic wells. Detection of p24_Gag_ with AlphaLISA was used to quantify the amount of captured virus in each well (Fig. 5A, top workflow). For this proof of principle test, we chose to use transfected viruses with or without PSGL-1 so that we could be certain that the differences we saw in P-selectin binding could be specifically attributed to the presence of PSGL-1. We observed high levels of virion capture in the P-selectin coated wells overlayed with PSGL-1^High^ virus, but not in P-selectin coated wells tested with control (PSGL-1^Neg^) virus. Uncoated wells also displayed minimal levels of non-specific capture (Fig. 5B).

**FIG 5.**
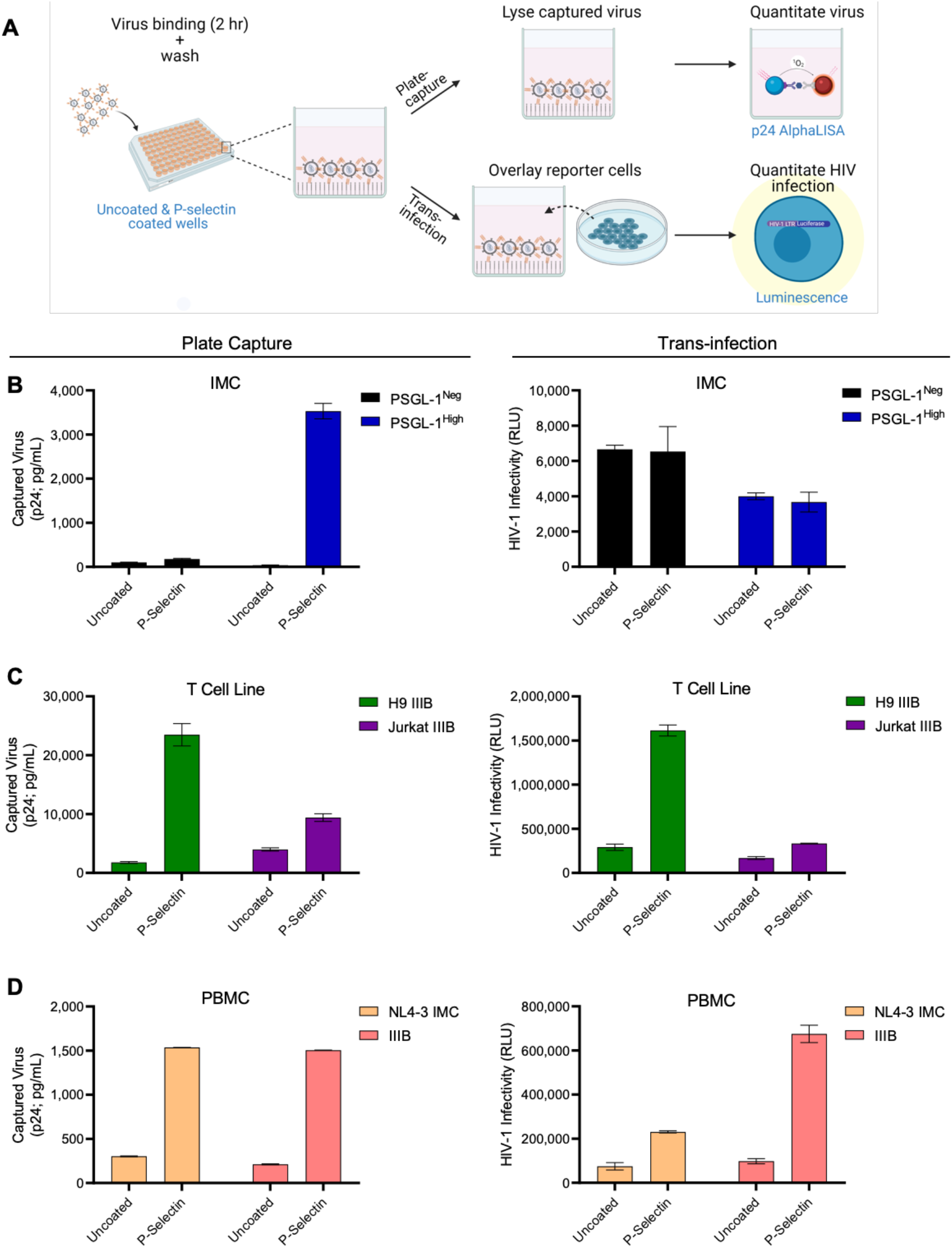
PSGL-1-positive virions can be captured by P-selectin and transferred to HIV-permissive target cells. (A) Schematic depicting the experimental workflow: Virus preparations were added to uncoated and P-selectin coated wells for two hours at room temperature to allow for binding of virion-incorporated PSGL-1 to P-selectin. Post-incubation, wells were washed extensively to remove unbound virus and then were either assayed for the amount of captured virus using p24 detection, or TZM-bl cells were overlayed onto each well, and infection was measured via luminescence. (B) Experimental results from IMC, (C) T cell line, and (D) PBMC viruses in plate-based virus capture (left column) and trans-infection assays (right column). Results are displayed as mean +/-standard deviation of samples tested in duplicate.

Since the primary target of HIV-1 infection, CD4+ T cells, are often found on activated endothelial tissues which display P-selectin in inflammatory conditions (54–56), we were interested in testing whether virions captured by P-selectin could be transferred to nearby permissive cells to facilitate trans-infection. We hypothesized that this could suggest new roles for virion-incorporated PSGL-1 beyond its current scope as an antiviral molecule. As a model to simulate trans-infection that could occur *in vivo*, we overlaid TZM-bl reporter cells on the wells containing P-selectin-captured virus (after washing away unbound virus) and assessed infection 48 hours later (Fig. 5A, bottom workflow). For these initial capture-transfer tests we chose to use replication-competent IMC viruses, rather than pseudoviruses, since we anticipated that multiple rounds of virus replication would be necessary to visualize detectable differences in infectivity caused by P-selectin-mediated capture, which would not be possible with pseudoviruses. As expected, PSGL-1^Neg^ and PSGL-1^High^ IMC viruses showed negligible levels of capture-transfer, with a level of background luminescence that is typical of uninfected cells, suggesting that no infection occurred (<8,000 RLU; Fig. 5B). These observations were in line with our expectations since minimal amounts of PSGL-1^Neg^ virus was captured using P-selectin and the PSGL-1^High^ viruses are not infectious (Fig. 1).

Having validated that PSGL-1 on transfected virions could be captured by P-selectin and that our model of trans-infection was specific and sensitive, we next decided to test this model with T cell line and PBMC viruses which contain lower levels of PSGL-1 and higher levels of gp120. We observed that viruses produced in both T cell lines and primary PBMC were captured by P-selectin and could be transferred to reporter cells for robust detection of infection (Fig. 5C and 5D). Virion capture with HIV-1 IIIB produced in Jurkat cells was present at much lower levels than those seen with the same isolate produced in H9 cells; however, this finding displayed a similar trend as the earlier anti-PSGL-1 antibody capture assays using these viruses (Fig 2), suggesting that the differences in virion capture may be related to differences in the two producer T cell lines. More importantly, both PBMC viruses were effectively captured by P-selectin and transferred to HIV-permissive cells, suggesting that this mechanism of capture-transfer may also be possible with P-selectin expressed on cells *in vivo*.

As an additional test to confirm that the plate-based virion capture seen using recombinant P-selectin was not simply caused by non-specific binding of virus to the adhesion protein, we tested the T cell line viruses in a similar capture-transfer assay whereby we substituted the capture molecule of P-selectin with antibodies specific for PSGL-1, gp120 or a non-specific isotype control (Fig S4). Remarkably, virions were captured with the anti-PSGL-1 antibody and transferred to permissive cells with similar efficiency as that seen with P-selectin capture (comparing data in Fig S4B to Fig. 5C). Also, the relative levels of plate-based capture between anti-PSGL-1 and anti-gp120 were similar to those previously observed in bead-based capture assays (Fig. 2). Taken together, these data provide three validations of our model system: 1) that the capture-transfer mediated by P-selectin capture is reproducible between different capture molecules for the same protein (anti-PSGL-1 vs. recombinant-P-selectin); 2) that our assay model can be reliably applied to capture with other virion-incorporated proteins (like gp120); and 3) that our trans-infection model system can reflect differences in the level of virion-incorporated proteins, as we observed less capture-transfer when capturing with anti-gp120, which is less abundant on virions.

## DISCUSSION

While PSGL-1 was first identified as a host restriction factor in a proteomic screen of activated primary T cells (10), many of the subsequent experiments characterizing the antiviral properties of the protein have been performed with cells transfected to ectopically express PSGL-1 (11–15). Although the bulk of evidence for the antiviral properties of PSGL-1 has been obtained using transfection models, as also confirmed in the present study, we show that PSGL-1 transfection models can overestimate the restriction effect of PSGL-1. Furthermore, these models are not always representative of biologically-relevant viruses produced by natural infection of T cells and primary PBMC. While endogenous levels of PSGL-1 on virions can also reduce infectivity (11, 12), when using transfection systems to study the impacts of PSGL-1 and other host molecules on viruses it is important for researchers to tailor transfection systems to resemble physiological conditions. An example of that is demonstrated herein using our PSGL-1^Low^ virus, which was the most similar to the more biologically relevant model systems (T cell line and PBMC viruses).

Our work shows that virions produced from T cells and PBMC remain infectious, with more virion-incorporated gp120, and markedly lower amounts of virion-incorporated PSGL-1 compared to transfected viruses which overexpress PSGL-1. Unique to this study, our observations were made with both a semi-quantitative virus capture technique which assessed the averages of the virus populations, and flow virometry, which can phenotype individual virions with high sensitivity (45). Our results showing low/undetectable levels of PSGL-1 on primary viruses, yet high levels of detection on transfected viruses, despite the use of orthogonal techniques, corroborates the notion that different virus production systems can lead to very different virus phenotypes. While the antiviral impact of PSGL-1 is clearly heightened when unnaturally high levels of PSGL-1 are present in the HIV-1 envelope, we demonstrated that transfected models can be generated with very low levels of PSGL-1 pDNA to be more representative of primary viruses, as seen with our PSGL-1^Low^ PV and IMC viruses. Indeed, initial work characterizing the antiviral properties of PSGL-1 utilized a wide range of pDNA (0-600 ng) to investigate the dose-response of infectivity in the presence of PSGL-1 (11, 15), which informed our selection of the range of pDNA values (0-1000 ng; Table S1) used to generate the low, medium and high PSGL-positive virions in this work. Of note, using calibrated flow virometry we were able to estimate that the number of PSGL-1 molecules on transfected pseudovirus preparations were as low as 15 and as high as 145 proteins per virion on the PSGL-1^Low^ and PSGL-1^High^ viruses, respectively. The values for the PSGL-1^High^ pseudoviruses are in a similar range to those which our group has reported previously (100 ± 57 SD; median ± SD) on PSGL-1 pseudoviruses generated without HIV-1 Env (28). While we acknowledge the limitations of the technique and caution that MESF values are estimations and not absolute quantifications, it is very likely that the number of PSGL-1 proteins on viruses produced in PBMC is less than 10-20 molecules per virion, since we were unable to detect the protein using flow virometry and we believe our limit of detection with this assay to be ∼10 MESF. Optimizing the threshold for detection of low abundance antigens, namely PSGL-1 and gp120, on virions produced in primary cells and patient samples using flow virometry remains the scope of future work to be performed by our group.

It is important to mention that while the addition of PSGL-1 to the HIV envelope has been shown to decrease levels of Env within virions *in vitro* (11, 15), all of the viruses produced in PBMC were shown to have detectable levels of Env, despite the presence of PSGL-1. Interestingly, this was not the case for transfected pseudoviruses, highlighting the need for careful selection of a virus production model in order to generate results that are physiologically relevant to viruses *in vivo*. Among the patient plasma samples assayed, PSGL-1 was found to be present on all of the viruses tested and at similar levels to the positive control, CD44. Previous work has shown that CD44 can efficiently capture viruses from plasma of HIV-infected patients (49, 50), and our results herein suggest that PSGL-1 could also be used for similar purposes in the future.

In this work we found that viruses propagated in T cell lines were much more similar to viruses passaged in primary cells than those produced through transfection of HEK293 cells. Although the antiviral role of PSGL-1 is known to be antagonized by the viral accessory proteins Vpu and Nef (10, 11, 15), it is possible that additional counteracting factors may be at play in viruses produced via infection of primary cells (and T cell lines). We hypothesize that this is the case since we observed that virion-incorporated PSGL-1 was much less abundant in viruses from these model systems compared to IMCs transfected in HEK293 cells, despite Vpu and Nef being present in the full-length IMC viruses. Though clear differences were apparent in the PSGL-1:gp120 ratios of transfected and PBMC viruses produced *in vitro*, the fact that PSGL-1 was incorporated in all of the PBMC-produced virus isolates tested, including SIV strains, demonstrates the biological importance of PSGL-1 in HIV infection and also supports the idea that PSGL-1 can play a role in the pathogenicity of a many enveloped viruses (11, 13, 14).

While the glycosylation of PSGL-1 is known to be cell-type dependent and critical for the ability of the protein to bind P-selectin (3, 4, 7, 22, 23), here we saw that PSGL-1 was capable of binding P-selectin on all of our model virions, regardless of the cell type used for virus production. This finding provides the basis for further studies to explore this role of PSGL-1 on virions within diverse tissue reservoirs, in *ex vivo* models and *in vivo* studies. Importantly, several other host proteins that have been shown to be virion-incorporated, such as ICAM-1 and integrin α4β7, are also known to maintain their functional activity and to be able to bind their cognate binding ligands and receptors, respectively (17, 24). Likewise, both of these proteins have been shown to alter or enhance HIV-1 pathogenesis in a humanized mouse model (24) or *ex vivo* human tissue (52, 57), respectively.

Since P-selectin is known to be expressed on endothelial tissues *in vivo* upon cellular activation (5, 58, 59) and is important in recruiting HIV-1 target cells such as CD4+ T cells into intestinal tissues (60), it is reasonable to speculate that PSGL-1 present on virions could also mediate binding and extravasation of virions into inflamed tissues, similar to what occurs with PSGL-1-expressing leukocytes (4, 61). Similarly, it is tempting to consider the possibility that HIV-1 virions may exploit incorporated-PSGL-1 to target intestinal CD4+ T cell populations and fuel local viral spread, similar to what was shown with integrin α4β7 in our previous work (24). This proposed function of PSGL-1 as a viral homing molecule would be especially relevant for HIV-1 pathogenicity during the early phases of infection when the gut is still highly populated with uninfected CD4+ T cells (62). However, this hypothesis remains untested and more targeted experiments with improved model systems of trans-infection that are more physiologically relevant will be the scope of future studies by our group. Regardless of these outcomes, it is important to consider that functional PSGL-1 on virions may have uncharacterized impacts *in vivo,* contrary to the predominant inhibitory properties that PSGL-1 was previously shown to exert *in vitro*.

Notably, in addition to being a ligand for P-Selectin, PSGL-1 can also bind E- and L-selectins depending on the protein’s glycosylation state (63–65). While the capability of virion-incorporated PSGL-1 to bind other selectins was not investigated here, this may be of interest in future work since this may provide insights into additional *in vivo* roles of this virion-incorporated protein.

Lastly, while this work contributes to a better understanding of the role of PSGL-1 in HIV-1 infection, this protein may also have uncharacterized impacts on other microbial pathogens. For example, PSGL-1 is known to be a functional entry receptor for enteroviruses (66) and is also known to be involved in *S. aureus* infection, wherby the bacterial fibrinogen inhibits P-selectin-PSGL-1 interactions *in vitro* (67). Our work opens the field for new paradigms on how PSGL-1 can impact microbial infections, and further extends the recently established paradigm that PSGL-1 is a potent host-restriction factor against HIV-1 viruses. The recent identification of PSGL-1 as a virion-incorporated restriction factor has fuelled a rapidly expanding field of literature. Our work is the first to identify dual roles for PSGL-1, in which it can both restrict and facilitate HIV infection, highlighting the need for continued work in this field. As a rapidly evolving and higly sophisticated pathogen, it is unsurprising that HIV would be able to exploit a host restriction factor to enhance its pathogenesis. Our work underscores the importance of studying virion-incorporated proteins using a more holistic approach, as the impact of enhanced pathogenesis and virus restriction afforded by PSGL-1 are likely not mutually exclusive. Future studies are warranted to establish the *in vivo* role of virion-incorporated PSGL-1 in virus homing and trans-infection, which could pave the way toward the development of novel antiviral therapies.

## METHODS

### Cell culture

The HEK293 and TZM-bl cell lines used to produce transfected viruses and readout HIV infection, respectively, were obtained from the NIH HIV Reagent Program (ARP; Cat#103 and 8129) and were maintained in complete media comprised of DMEM (Wisent, Cat#319-005-CL), 10% fetal bovine serum (FBS; Wisent, Cat#098150), 100 U/mL penicillin, and 100 µg/mL streptomycin (Life Technologies, Cat#15140122). The H9 and Jurkat (E6-1) T cell lines (ARP Cat#87 and 177) and peripheral blood mononuclear cells (PBMC) used to produce virus through infection were maintained in RPMI-1640 (Wisent, Cat#350-000-CL) with the same components listed above. Recombinant human IL-2 (25 U/mL) was also added to PBMC cultures. All cells were grown in a 5% CO_2_ humidified incubator at 37 °C.

### Infection-based virus production

Whole blood from healthy donors was acquired in accordance with University of Toronto’s Research Ethics Board approval (protocol #00037384), with all donors providing written informed consent. Blood was collected in heparinized vacutainers (BD Biosciences) and PBMCs were subsquently isolated using density centrifugation with Lymphoprep (StemCell Technologies, Cat#07861). PBMCs were activated with 1% phytohemagglutinin (Gibco, Cat#LS10576015) and 50 U/mL of IL-2 in complete RPMI media for 72 hours prior to infection with primary HIV isolates. H9, Jurkat cells or activated PBMC were pelleted and resuspended in 1 mL of replication-competent HIV isolates BaL, NL4-3 IMC or IIIB for 4 hours to facilitate infection before fresh media was added to the cells. Cell culture supernatants containing virus were harvested 7-12 days later based on viral titre.

### Transfection-based virus production

HIV pseudoviruses and IMCs were produced with 2 µg of SG3^ΔEnv^ or pNL4-3 pDNA respectively and various levels of PSGL-1 and HIV-1 BaL.01 Env pDNA (as outlined in Table S1) to generate PSGL-1 negative, low, medium or high viruses (PSGL-1^Neg^, PSGL-1^Low^, PSGL-1^Med^, PSGL-1^High^). Viruses were produced in HEK293 using Polyjet *In Vitro* Transfection Reagent (FroggaBio, Cat#SL100688). HEK293 cells were seeded at a density of 10^6^ cells/mL in 6-well plates in complete media and were transfected with 3 µg of pDNA after cells had reached 70% confluence. All transfections were performed with a 1:3 ratio of plasmid DNA (µg) to transfection reagent (µL). Six hours after transfection, the medium was replaced with complete DMEM to discard any viral progeny without incorporated host proteins. Cell culture supernatants containing virus were harvested 48 hours after transfection. All HIV-1 expression vectors were acquired from the NIH ARP. The negative control and PSGL-1 expression vectors were obtained from Sino Biological (Cat#CV011 and HG10490-UT).

### Virus titration

For infection, TZM-bl cells were seeded into 96-well flat-bottom plates at 15,000 cells/well in 100 µL of complete DMEM. Virus stocks were added to the cells, yielding a total volume of 200 µL/well. Viral input was normalized by p24 for each virus cluster tested (except for PBMCs which were run at their undiluted viral titre due to the wide range of variation between donors). Luciferase expression was detected 2 days later after removal of 135 µL of medium and the addition of 45 µL per well of Britelite™ Plus reporter assay (PerkinElmer, Cat#6066766). The cell lysates were transferred to Perkin Elmer Optiplates after 10 minutes for luminescence endpoint readout on a Synergy neo2 (BioTek) microplate reader. All the samples were tested in duplicate.

### Antibodies

The following mouse and human monoclonal antibodies were utilized for virion capture assays: anti-PSGL-1 and anti-mouse IgG (BD Biosciences, Cat# 556053 and 557273) and anti-gp120s (clones 2G12 and PG9 acquired from the NIH ARP; Cat#1476 and 12149). For virion capture assays in patient plasmas, we used biotinylated antibodies, including CD44-bio (Miltenyi Cat#130-113-340) and an in-house biotin conjugation (Bio-Rad Laboratories, Cat#LNK262B) of anti-PSGL-1 and mouse IgG1 isotype control (R&D Cat#MAB002) antibodies. R-phycoerythrin (PE) conjugated goat anti-human IgG Fc secondary antibody (Invitrogen, Cat#12-4998-82) and PE conjugated mouse anti-PSGL-1 mAb (BD Biosciences, Cat#556055) were used for flow virometry labelling.

### Flow virometry

Flow virometry was performed using a Beckman Coulter CytoFLEX S with standard optical configuration and volumetric calibrations were performed as described previously (28). Gain and threshold optimization for detection of virus and calibration beads was performed as described previously (68), with a modification of PE gain to 2600(68)(69). All virus samples and controls were acquired at a sample flow rate of 10 µL/min for 1-2 min. Serial dilutions of select stained viruses were acquired on the cytometer to determine the presence of viral aggregates and to control for coincidence (Suppl File 2). For direct labelling, cell-free supernatants containing virus were diluted to 10^9^ particles/mL, or used undiluted if they were estimated to be less concentrated based on viral p24 data and stained overnight at 4°C with a PE-conjugated mAb against PSGL-1. For indirect antibody labelling, an unconjugated PSGL-1 mAb of the same clone was used with the secondary antibody described above on similarly diluted virus samples. After labelling, all samples were further diluted with PBS (to reduce coincidence) for analysis by FV. Virus particle concentrations were determined by flow cytometry by gating on the virus population using SSC (Fig S1). After staining virus stocks were diluted two-fold with 4% PFA (2% final) for 30 minutes for fixation. BD Quantibrite PE beads (CA, USA; Cat#340495, lot 91367) and Spherotech 8 Peak Rainbow calibration particles (Cat# RCP-30-5A, lot AF01) were used for fluorescence calibration, while NIST-traceable size standards (Thermo Fisher Scientific) were used for light scattering calibration (see Suppl File 2 for full list). Light scatter calibration was performed using FCM_PASS_ software (https://nano.ccr.cancer.gov/fcmpass) as previously described (68, 69). Cross-calibration of fluorescence was also performed using FCM_PASS_ for the PE channel as described elsewhere (69). Detailed information on the fluorescent and light scatter calibration and the MIFlowCyt-EV checklist (70) can be found in the FCM_PASS_ output report in Suppl File 2 along with the FCM_PASS_ data plots. All data were analyzed using FlowJo software version 10.7.1. (CA, USA). The median and robust standard deviation statistics for PE MESF were generated using the strategy described in Figure 3 using FlowJo.

### Virion capture assay

Immunomagnetic bead-based virion capture was performed as previously described (24, 35), with 25 µL of protein G Dynabeads (Life Technologies; Cat#10004D), which were armed with 0.5-1 µg of anti-PSGL-1 or anti-gp120 for 30 min at room temperature and then washed with 10% FBS–PBS to remove unbound antibodies. Where indicated the virus input was normalized at the same concentration (35 ng/mL of p24) across all viruses tested. For all tests equal virus volumes were used. Viruses were incubated with antibody-armed beads for 1–2 h at room temperature to allow virus capture. Beads were then washed twice with 10% FBS–PBS and once with 0.02% FBS–PBS to extensively remove unbound virus particles. The bead-associated virus was then treated with 0.5% Triton X-100 to lyse the captured virions for p24 quantification by AlphaLISA. Data analysis was performed using Prism v. 8.4.2 (GraphPad, San Diego, CA, USA). The background level of virion capture for each virus type was assessed by virion capture with an isotype control antibody (as described above). The nominal level of background capture as detected using the non-specific isotype antibody was removed from each data point before graphing where indicated.

For experiments assessing viruses in plasma of HIV-infected patients, biotinylated antibodies (described above) were used to permit more sensitive virus capture assays with anti-biotin microbeads (Miltenyi, Cat#130-090-485). Levels of captured virus were measured by HIV-1 RNA detection via quantitative real-time PCR assays. Viral RNA was purified using the QIAamp Viral RNA kit (Qiagen) and the number of HIV-1 genome equivalents was obtained using previously reported primers, probe, and amplification conditions (71). Plasma samples (patient ID #’s 3-5 and 8-12) were contributed by Drs. Tae-Wook Chun and Susan Moir, collected in accordance with a protocol approved by the Institutional Review Board of the National Institute of Allergy and Infectious Diseases, National Institutes of Health (ClinicalTrials.gov number NCT00039689). Dr. Frank Maldarelli provided additional clinical specimens (ID #’s 1-2 and 6-7) under approved NIH protocols. All subjects provided written informed consent.

### p24 AlphaLISA

The quantification of HIV-1 p24 capsid protein was performed in captured virus lysates and virus-containing supernatants with the high-sensitivity AlphaLISA p24 detection kit following the manufacturer’s (PerkinElmer) instructions. Absorbance readings were performed on a Synergy NEO 2 multimode plate reader (BioTek, VT, USA) equipped with Gen 5 software (v. 3.08).

### Plate capture and trans-infection assay

Sterile tissue culture plates were coated overnight at 4°C with 5 µg/mL recombinant CD62P protein (R&D Systems, #137-PS-050). The following day the wells were washed 3X with PBS before being blocked with 1% BSA in PBS at room temperature (RT) for 1 hour. Following blocking, the wells were washed as before, and virus stocks were added to the wells for 2 hours at RT. After viral incubation, the wells were washed to remove unbound virus and were either treated with 0.5% triton X-100 in PBS to lyse captured virus or 15,000 TZM-bl cells were added to the wells to detect the ability of captured virus to be transferred to permissive cells. The viral lysates were assayed for p24 and TZM-bl cell luminescence was read 1-4 days later depending on the starting concentration of virus used.

## SUPPLEMENTAL MATERIAL

**SUPPLEMENTAL FILE 1:** Table S1, Figure S1, Figure S2, Figure S3, Table S2, Table S3, Figure S4.

**SUPPLEMENTARY FILE 2:** Flow virometry FCM_PASS_ output report, with fluorescent and light scatter calibration, and MIFlowCyt-EV checklist.

## ACKNOWLEDGMENTS

We thank Drs. Tae-Wook Chun, Susan Moir, and Frank Maldarelli for providing the plasma samples from HIV-infected individuals used in this study. The following reagents were obtained through the NIH HIV Reagent Program, Division of AIDS, NIAID, NIH: Jurkat E6-1 cells (ARP-177), H9 cells (ARP-87), TZM-bl cells (ARP-8129), HIV-1 SG3^ΔEnv^ non-infectious molecular clone (ARP-11051), HEK293 cells (ARP-103), HEK293 cells (ARP-103), pNL4-3 (ARP-3417), 2G12 mAb (ARP-1476), PG9 mAb (ARP-12149) and HIV-1 BaL.01 Envelope expression vector (ARP-11445), respectively contributed by ATCC (Dr. Arthur Weiss), Dr. Robert Gallo, Dr. John. C. Kappes and Dr. Xiaowun Wu (generously donated both TZM-bl and SG3^ΔEnv^ non-infectious molecular clone), Dr. Andrew Rice, Dr. Nathaniel Landau, DAIDS/NIAID, International AIDS Vaccine Initiative, and Dr. John Mascola. The authors acknowledge the University of Ottawa Flow Cytometry and Virometry Core Facility for the acquisition of the flow virometry data. All flow virometry data calibrations were completed using FCM_Pass_. All illustrations were created with BioRender. This research was supported in part by the Intramural Research Program of the NIAID, NIH.

## Supplemental Legends

**Table S1. Amounts of plasmid DNA used to produce the BaL pseudovirus (PV) and NL4-3 infectious molecular clone (IMC) viral stocks.** Virus stocks with different amounts of virion-incorporated PSGL-1 were created (PSGL-1 negative, low, medium, or high). Empty vector plasmid was included in the IMC virus transfection to ensure an even amount of total pDNA (3 µg) was transfected in all conditions, irrespective of the variable amount of PSGL-1 plasmid used. The PV transfections also included an even amount of pDNA (3 µg) across all conditions, with minor fluctuations in Env pDNA to account for the differential amounts of PSGL-1 pDNA used.

**FIG S1. Virus populations can be distinguished from background levels of fluorescence present when acquiring culture media.** Unstained cell culture medium alone (media) or cell culture supernatants containing transfected HIV (PSGL-1^High^ PV, media + virus) are shown with gating on the virus population. Particle concentrations of the bottom gated regions are shown in red on each dot plot. Lower gates are set on the side scatter population of viruses as described previously (PMID: 33198254). The lower gate spans 10-60 nm² on the x-axis and has an upper limit of 10 PE MESF on the y-axis.

**FIG S2. Titration of anti-PSGL-1 PE-conjugated antibody on all virus stocks.** (A) Titration of anti-PSGL-1-PE antibody on pseudoviruses and (B) IMC viruses produced via transfection of HEK293 cells. Cell culture supernatants containing virus or cell culture medium alone were stained overnight at 4°C with four antibody concentrations (as indicated) or acquired unstained on the cytometer. (C) Staining and acquisition were performed as in (A) on IIIB viruses produced through infection of the H9 and Jurkat T cell lines or (D) BaL, IIIB or NL4-3 IMC viruses produced in activated PBMCs. Events above the dashed lines indicate positive PSGL-1 labelling. This line was set directly above the level of background fluorescence seen on unstained virus or cell culture media (DMEM for transfected viruses and RPMI for viruses produced through infection).

**FIG S3. Comparison of staining PSGL-1-positive viruses with one-step versus two-step staining procedures.** (A) Staining of PSGL-1 on transfected viruses with (PSGL-1^High^) and without (PSGL-1^Neg^) PSGL-1 in the viral envelope using either an anti-PSGL-1 PE-conjugated primary antibody (upper panel) or an unlabelled mouse anti-PSGL-1 antibody paired with a PE-labelled anti-mouse secondary antibody (lower panel). (B) A comparison of PSGL-1 staining on both virus populations in units of PE MESF are shown in red, as identified from the population of gated viruses that appear above the horizontal dotted line (>10 PE MESF on the y-axis) on virus dot plots from (A). This line denotes the limit of instrument detection. An equivalent vertical line depicted on each histogram denotes the same level of background fluorescence. PSGL-1 staining on control viruses (PSGL-1^Neg^) that yielded MESF values below 10 are not displayed.

**Table S2. Additional clinical information on HIV-infected patients from which plasmas were derived.**

* Patient ID symbols correspond to patient data points shown in Fig. 4.

† Acute/early infection is estimated to be less than 6 months from the putative infection date.

**Table S3. Percentage of total virus captured using anti-IgG, -PSGL-1, or -CD44 antibody capture from viremic plasmas (from Figure 4).**

**FIG S4. Method validation for plate-based virus capture and trans-infection assay.** (A) Schematic depicting the experimental workflow: Virus preparations were added to wells precoated with mAbs specific for either PSGL-1 or gp120, or with an isotype control (IgG) for two hours at room temperature to allow for virus binding. Post-incubation, wells were washed extensively to remove unbound virus and then were either assayed for the amount of captured virus using p24 detection, or TZM-bl cells were overlayed onto each well containing captured virus, and infection was measured via luminescence. (B) Experimental results from plate-based antibody-mediated virus capture (left panel) and trans-infection assays (right panel). Results are displayed as mean +/-standard deviation of samples tested in duplicate.

**Supplementary Data File 2: FCM_PASS_ outputs from fcs file calibration.**

